# Controlling fluid flow to improve cell seeding uniformity

**DOI:** 10.1101/299966

**Authors:** Paul M. Reynolds, Camilla Holzmann Rasmussen, Mathias Hansson, Martin Dufva, Mathis O. Riehle, Nikolaj Gadegaard

## Abstract

Standard methods for seeding monolayer cell cultures in a multiwell plate or dish do not uniformly distribute cells on the surface. With traditional methods, users find aggregation around the circumference, in the centre, or a combination of the two. This variation is introduced due to the macro scale flow of the cell seeding suspension, and movement of the dish before cells can settle and attach to the surface. Reproducibility between labs, users, and experiments is hampered by this variability in cell seeding. We present a simple method for uniform and user-independent cell seeding using an easily produced uniform cell seeder (UCS) device. This allows precise control of cell density in a reproducible manner. By containing the cell seeding suspension in a defined volume above the culture surface with the UCS, fluctuations in cell density are minimised. Seeding accuracy, as defined by the actual cell density versus the target seeding density is improved dramatically across users with various levels of expertise. We go on to demonstrate the impact of local variation in cell density on the lineage commitment of human embryonic stem cells (hESCs) towards pancreatic endoderm (PE). Variations in the differentiation profile of cells across a culture well closely mirror variations in cell density introduced by seeding method – with the UCS correcting variations in differentiation efficiency. The UCS device provides a simple and reproducible method for uniform seeding across multiple culture systems.

## Introduction

Experiments involving cell culture, from biomaterial testing^1–3^ to drug discovery^4^ often begin with cells seeded onto a flat surface to form a two dimensional culture. This is the foundation on which the experiment as a whole is built and is arguably one of the most critical steps. Seeding density influences cellular behaviour in sparse versus dense cultures due to differences in cell-cell communication, local concentration of auto- and paracrine factors, cell shape and mechanical interaction. The commitment and differentiation of stem cells are highly regulated by cell density, and so the initial seeding density has been shown to influence the differentiated phenotype of pig articular chondrocytes in culture^5^, human embryonic stem cell (hESC) differentiation towards pancreatic endocrine cells^6^, and other cell fate decisions^7,8^. Issues with uneven seeding also arise in the case of array and screening platforms, whereby a disparity in cell density can introduce noise and variability – leaving the assays open to errors^6,9–11^. Whilst rarely considered in the literature, uneven seeding may also skew results when conducting biomolecular assays across an entire well, including measures in supernatant, cell lysate and DNA/RNA^12^. Human bone marrow cells plated at high density show increased Notch signalling^13^, density dependent metabolic profiles^14^, and modified viability^15^ – which are measurements aggregated from a single culture vessel with uneven cell density.

Uneven seeding arises due to three factors – the macro scale turbulent flow of cell seeding suspension as it is added to the dish or well, disturbing the cell suspension as plates are moved to the incubator, and to a lesser extent the meniscus that forms around the wall of the culture plate (this effect is more pronounced with smaller well sizes). The problems associated with uneven cell seeding are apparent in the volume of forum posts^16^ from students seeking help in improving their seeding, after struggling with variability. Studies investigating the optimal means of cell seeding have sought to identify the source of uneven cell distribution^17^ but the number of published studies lags behind the apparent need expressed online. There are commercial culture vessels available which remove the meniscus effect^18^, purporting to improve uniformity in hematopoietic CFU assays - but they still suffer from uneven distribution due to the remaining effects of fluid flow. Common practice for this crucial step also varies between disciplines, laboratories, individual researchers, and even from day to day. Furthermore, high user dependence of cell seeding introduces variability to the results and leaves others struggling to reproduce work^19–21^. Suggested protocols exist that are not uniformly adopted, nor included in the detailed methods as they are deemed too basic to mention^22^. These standard laboratory methods additionally still cause variations in density across the vessel^23^. Applying our method effectively eliminates this variation. In light of this, achieving a consistent and reproducible seeding density across each sample is imperative for the production of high quality data^14^.

## Results

### A simple device to increase seeding uniformity

Redistribution of cells in an uneven manner can occur due to fluid flow and vessel movement after seeding (**Fig. 1A**). We propose a method of cell seeding which makes use of a simple lid above the culture surface to confine the cell suspension and maintain uniform seeding volume per unit area. This uniform cell seeder (UCS) device has only two strict design parameters: 1) the upper surface is flat and rigid, and 2) it is close enough to the target surface to efficiently wick the suspension into the cavity and hold it by capillary force (**Fig. 1B**). Rather than dispensing a droplet of cell suspension onto a sample, or pipetting cell suspension over a surface already immersed in media, the UCS is placed on the surface and cell suspension is pipetted through an inlet (**Fig. 1C** & **Video S1**). After moving the culture vessel to an incubator to allow for cell attachment (2-4h) the vessel is back-filled with culture media. The UCS device floats away from the surface and can be removed without disturbing the culture. The device corrects for seeding artefacts such as clustered regions of high and low density, however some edge effects remain in the seeding of well plates due to the meniscus formed by the cell suspension at the outer circumference (**Fig. 1D**).

**Fig 1.**
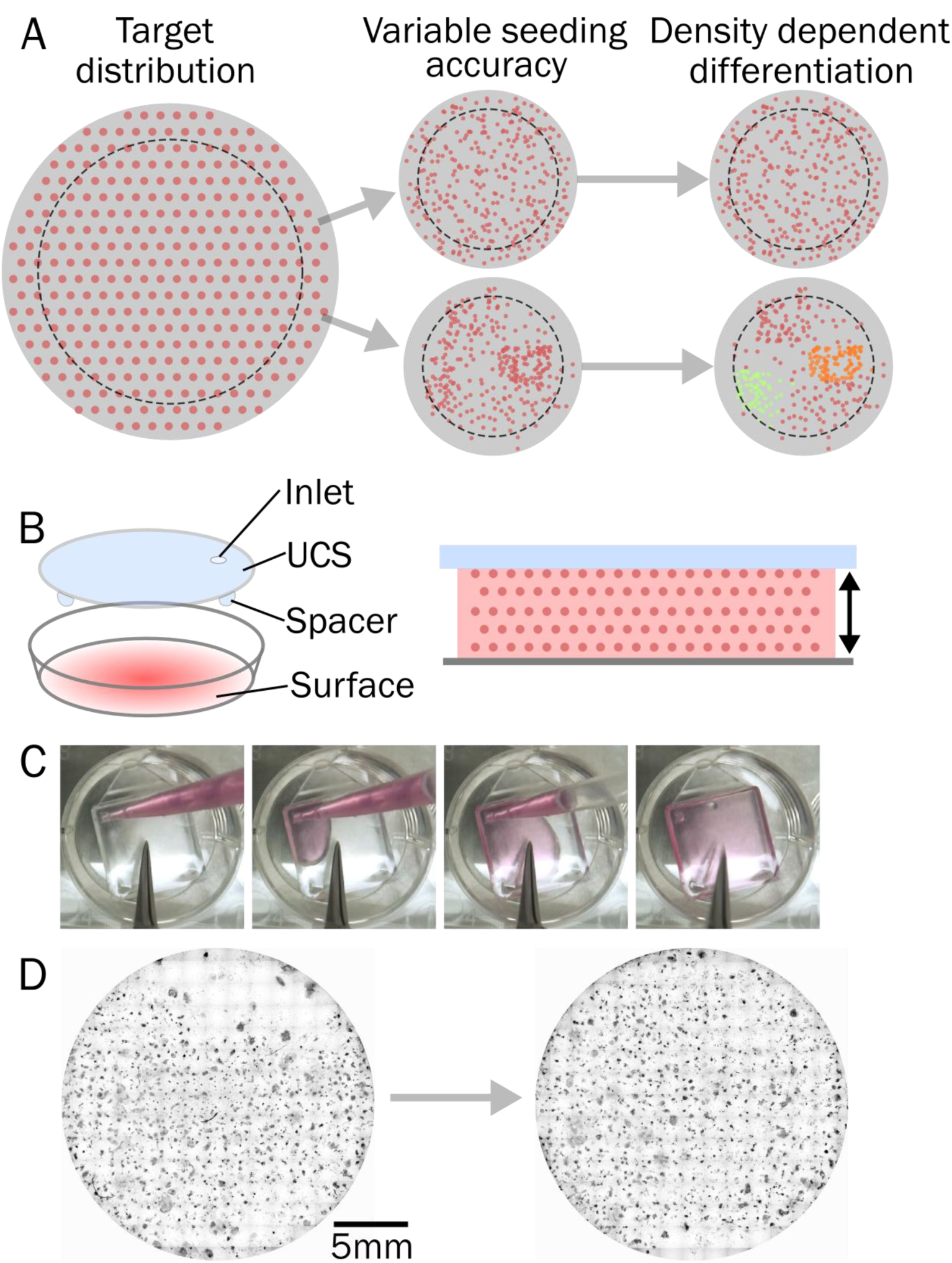
Improving the uniformity of cell seeding. **(A)** Uniform spacing between each cell (red dots) is the targeted distribution of cells in a dish when seeding to form a monolayer. The reality is that the same number of cells aggregate around the well edge and well centre resulting in a heterogeneous density distribution. Dashed outline shows example of a surface such as a coverslip in the well, which would often not be centred and suffer from further uneven distribution. Movement of the culture vessel to the incubator causes further redistribution of cells due to fluid flow, ultimately resulting in variation in cell phenotype depending on local changes in density. **(B)** The uniform cell seeder (UCS) device can be used to control the cell suspension, and therefore cell density, across a surface. A capping lid (blue) above the culture surface (grey) holds the cell suspension (pink) at a uniform height across the area (arrows), which evenly distributes the cell population. **(C)** The UCS device is filled by pipetting the cell suspension through an inlet. In this case, a 22mm coverslip in a 24 well plate is seeded with a uniform distribution of cells. Through capillary force, the cell suspension is held between the UCS and culture surface. **(D)** Nuclear staining of cells seeded in a 24-well plate without (left) and with (right) the UCS device after 2 weeks culture. Some aggregation around the edges remains, but the uniformity across the well is significantly improved.

As the only design criteria is that the surface and lid are held at a constant distance, these devices can be fabricated by simple spacers attached to flat material which can be laser cut to custom sizes and sterilised in ethanol, by UV, or by autoclave. We have also demonstrated production by 3D printing and mass production of thousands of disposable devices by injection moulding (**Fig. S1**). To determine time required to allow settlement of a seeding cell suspension to the surface, settle time was measured on a modified internal reflection microscopy setup (**Fig S2**). Cells were introduced into the UCS device and imaged from below, only being illuminated when they contacted the surface. After 3 mins the majority of cells had settled to the surface, and after 10 mins the number of cells at the surface was in an established steady state. After this waiting time, the culture can be moved to the incubator with minimal disruption to the distribution.

## UCS improves seeding accuracy and uniformity in a group of users

To compare the proposed method against current techniques we seeded human fibroblast cells with regular and UCS controlled seeding. Six users with varying personal technique and levels of experience were tasked with seeding at three different densities in a 12-well plate, in as uniform a manner as possible. A cell suspension was provided for each density, and the multiwell plate for each user was carefully moved to the incubator after allowing cells to attach to the well for 10 mins at room temperature. After 24h, cells were fixed and stained with nuclear stain DAPI followed by imaging the full well area. Our results demonstrate a marked variation in uniformity, both between methods and also between users (**Fig. 2A)**. Plotting the cell number across the well showed aggregation in the centre at low and medium cell densities, and more uniform seeding at higher densities (**Fig. 2A**). This varied between users and was not necessarily improved by more experienced users. An increase in cell number at the well edge was more apparent for some users. Seeding at the same densities using the UCS to control the cell suspension effectively corrected seeding defects in all users at all tested densities– with similar edge effects across the user group (**Fig. 2A**). To quantify this improvement in seeding accuracy we calculate the mean absolute error (MAE) of observed cell density across the full culture area versus the target density (**Fig. 2B&C**). Given ideal seeding in which cells are perfectly distributed at the target density MAE = 0. We found MAE was reduced by a factor of 3 when the UCS device was used to seed the culture. Whilst the error was not completely removed by the UCS device, it modified the distribution of cell densities on the sample to normally distributed across users. This is as expected of the biological reality of cell seeding - wherein a number of cells will fail to attach and others will be clumped together.

**Fig 2.**
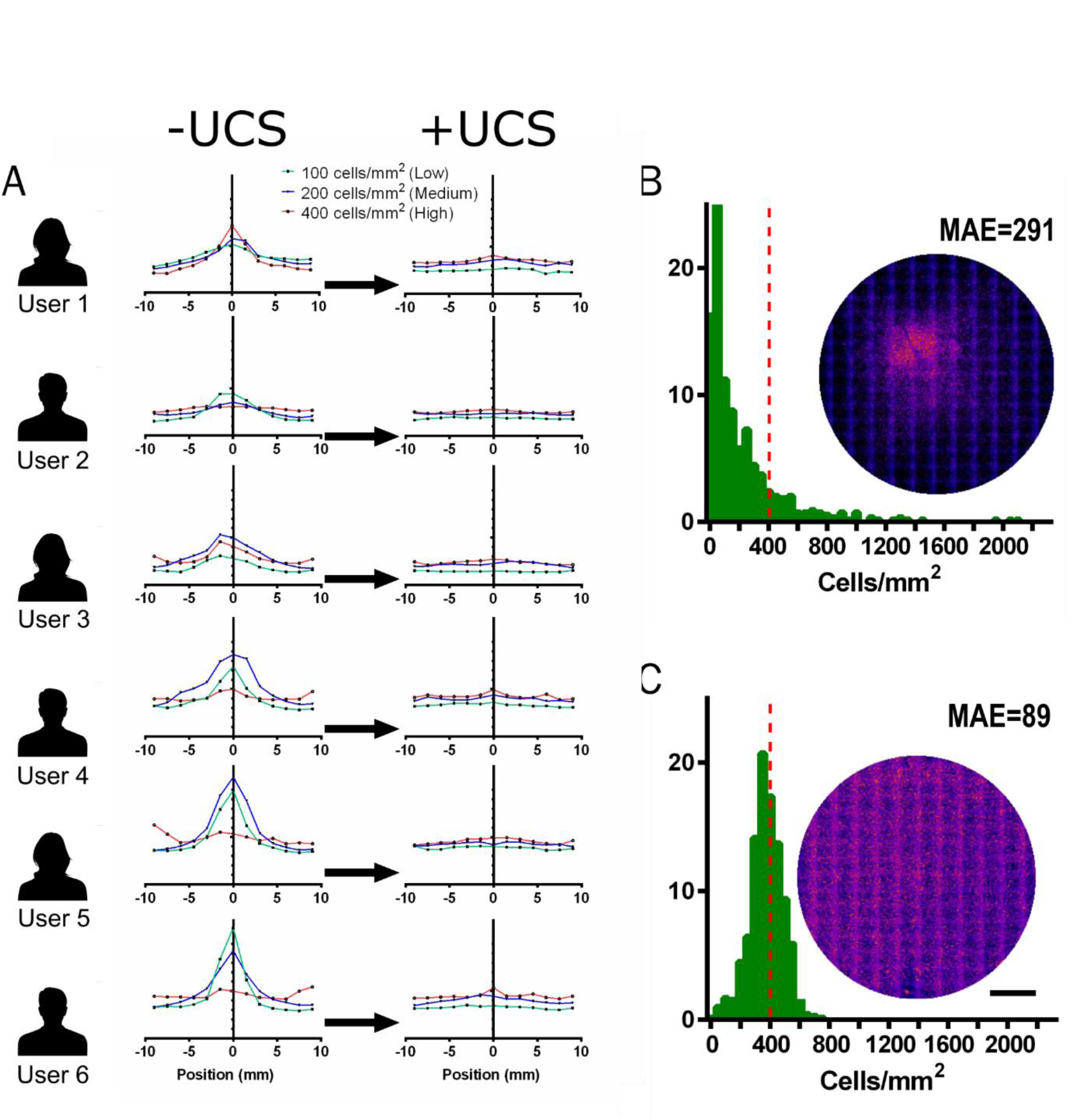
The UCS corrects seeding uniformity. (A) Six users of varying experience were tasked with seeding cells in a 12-well plate as evenly as possible at high (400 cells/mm^2^), medium (200 cells/mm^2^), and low (400 cells/mm^2^) densities. Cell number across the well centre is shown for each user without (left) and with (right) the UCS device. The y-axis shows the cell number normalised to the target density for each dataset indicated as a dashed line. (B) & (C) Histograms show the local cell density for all users (n=6) across the entire well after seeding with a target density of 400 cells/mm^2^ (indicated by red dashed line). Insets show heatmaps of n=6 samples seeded by each method. Mean Absolute Error (MAE) is shown for each method. Scale bar: 5mm

### Density dependent differentiation of hESC to the pancreatic endoderm

To demonstrate the impact of local density variations on the lineage commitment of stem cells, hESCs were seeded by standard methods and with the UCS device. We obtained similar results as with fibroblast cells in terms of seeding uniformity. Using the UCS we were able to obtain much more uniform seeding density as compared to standard seeding at all tested seeding densities (1000-3000 cells/mm^2^). We found that differentiation towards pancreatic endoderm (PE), as indicated by positive stain for PDX1, across the well depended on the seeding method and seeding density (**Fig. 3A-C**). With standard seeding of a low density (1000 cells/mm^2^), PDX1 positive cells were only observed in the centre of the well. Thus, differentiation into PE cells was only possible in areas with high cell density - which standard seeding provides only in the well centre while the UCS device mitigates. With intermediate seeding density (2000 cells/mm^2^) it was possible to obtain PDX1 positive cells both with standard seeding and UCS device seeding. However, a more uniform distribution of clusters of PDX1 positive cells across the entire well was obtained when using the controlled seeding technique compared to using standard seeding (**Fig. 3D**). With high seeding density (3000 cells/mm^2^) the differences in the seeding were less pronounced between the two different seeding techniques, however a change in PDX1 positive cluster distribution was still observed. It has been shown that patterning of hESCs into size controlled clusters can benefit yield of PE cells^24^ and we propose that the UCS is a simple means of repeatedly doing so. Variations in PDX1 expression across the well persist using the UCS device, but are far less pronounced. We assume that edge effects are caused by the local density of signalling factors dominating towards the well edge after backfilling the well with media, leading to a reduction in effective differentiation.

**Fig 3.**
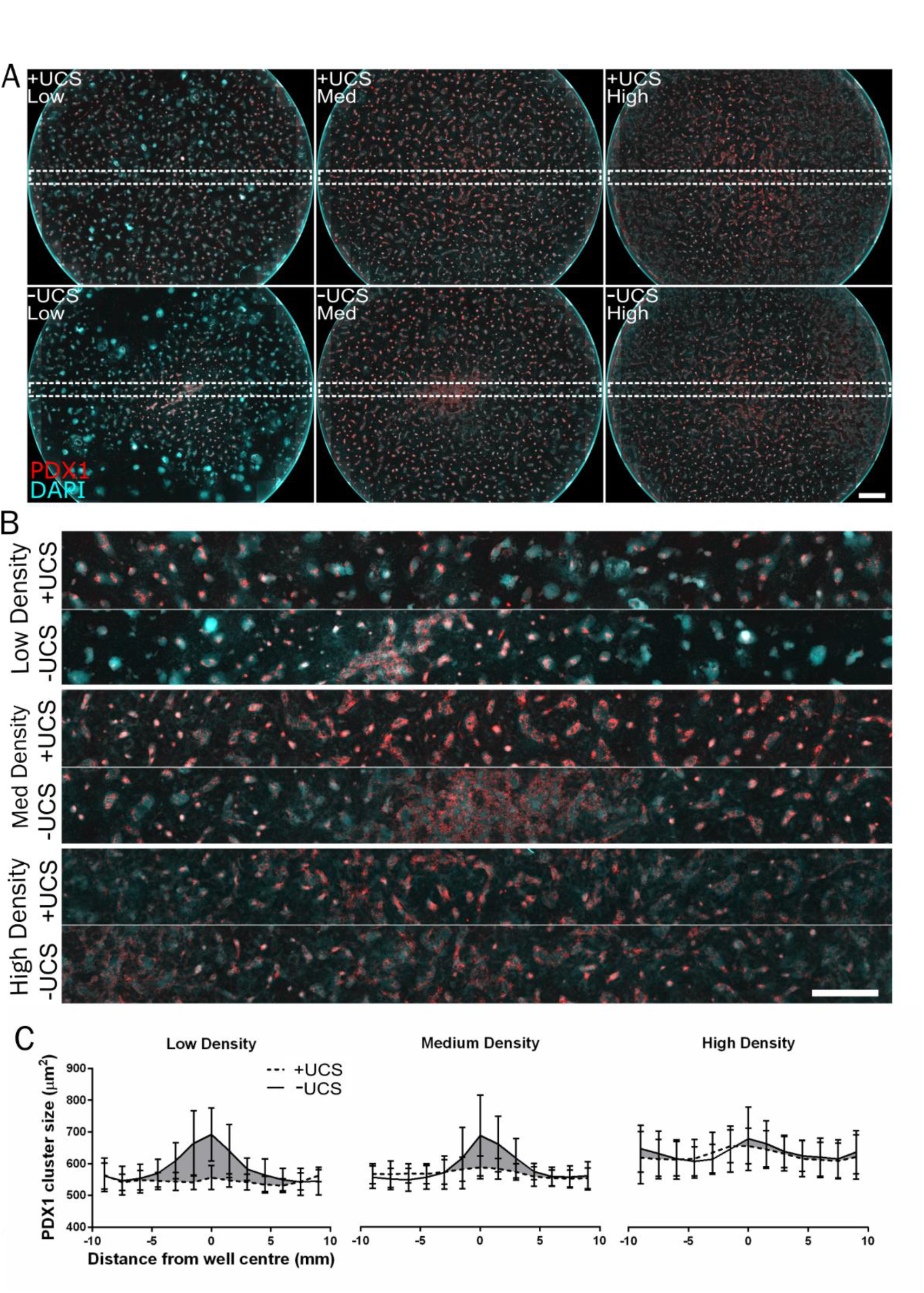
hESC differentiation mirrors seeding density across a well and can be controlled by the UCS device. **(A**) hESCs seeded with (top) and without (bottom) the UCS device at three seeding densities – low (1000 cells/mm^2^), medium (2000 cells/mm^2^) and high (3000 cells/mm^2^). A full 15×15 image array of a 12-well plate diameter is shown. PDX1 positive cells (red) and DAPI (blue). Scale bar: 2mm. **(B**) cross section montage of PDX1 labelled cells for each condition, as outlined in dashed white box in **(A)**. Seeding cells without the UCS device (labelled -UCS) results in central regions with higher expression of PDX1 whereas PDX1 expression is more evenly distributed where cells were seeded in an even manner. Scale bars: A = 1mm, B = 0.5mm. **(D**) Quantification of total PDX1 cluster area across the centre of the culture well, measuring average size of PDX1 positive clusters in across the centre (n=6, bars 95% CI).

## Discussion

In this study we present a simple to use device which effectively eliminates large variations in cell density across a culture vessel. Variability in the uniformity of cell seeding in a dish is a fundamental problem which is especially critical in studies of population sensitive cells. These cells respond to trophic, mechanical, and chemical factors presented by their local environment – and so changes in the local cell number across a culture vessel can result in entirely different cell phenotypes. By using a simple device to allow users to control the way in which cell suspension flows and covers the culture surface, we have removed variable aspects of cell seeding arising from fluid flow in the dish redistributing cells. The UCS corrects for differences in user technique and experience, to provide a repeatable cell seeding distribution without regions of high and low density in a single well. Consistent seeding at target cell densities improved reliability and reproducibility across experiments and users.

Monolayer cultures are seeded by one of two methods, either a droplet of fluid is placed on the surface (for example a glass coverslip), and the vessel back filled with culture media after cells have attached, or the coverslip is immersed in culture media, with a concentrated cell suspension then pipetted over the dish in as even a manner as possible. This is followed in some cases by a slight shake of the dish north/south/east/west to redistribute the cells evenly. Neither of these methods guarantee reproducible distribution of the cells, but are often considered adequate. The literature also suffers from confusion stemming from interchangeable use of seeding densities quoted in either cells/ml or cells/cm^2^. Methods to control and correct for uneven cell seeding in 3D cultures^25^ and microfluidic systems^26^ have been reported, however there are no published methods for improving uniformity in the most common culture vessels. The UCS device is a simple to fabricate tool which guarantees reproducible seeding by accurately controlling the distribution of cell suspension over the surface. It also lends itself to the reporting of cell seeding densities per unit area (i.e. cells/cm^2^) which can be easily reproduced between labs and users.

That variations in cell density occur after seeding, and that this variation in density is mirrored by variation in differentiation behaviour is not a surprising observation. It is well known that local cell density in the microenvironment plays a key role in determining cell fate and differentiation^6,23,27^. That such a simple device can provide such a drastic increase in uniformity may provide an important tool in improving the reproducibility of experiments. The extent of the problems associated with uneven cell seeding is not clear from the literature. Using the method described here we hope the community will assay their seeding uniformity, and attempt to understand how experimental procedures could be improved by controlled seeding with the UCS. Uneven cell seeding may even be advantageous for certain cell lines at low density, where valuable primary or pluripotent cells are used at the minimum acceptable concentrations. We saw increased expression of early PE markers in very low density cultures which were seeded in an uneven manner which was effectively removed by uniform seeding (**Fig. S5**).

The profile of cell density across multiwell plates seeded in **Fig 2** was mirrored in the profile of PDX1 positive cell cluster size after differentiation of hESCs to the pancreatic endoderm. The efficiency of differentiation was linked cell density, and in instances where cell density was not uniform across the plate at the time of seeding, it followed that differentiation was not uniform. The negative effects of cell seeding uniformity on differentiation consistency were not uniform across all densities in this case, with lower densities appearing to be more susceptible to larger differences in differentiation. Presumably, this is because regions of high local density passed a threshold for cell-cell signalling which induced differentiation more effectively. Understanding that the local cell distribution affects the differentiation potential of hESC towards their target differentiation state allows users to limit cells which are seeded and ultimately unlikely to differentiate. Protocols which appear to work, and use a large number of cells in a 12 or 24 well plate, may be more effective if used in a smaller culture area at higher density with uniform seeding.

By controlling the shape of the cell seeding suspension, the UCS device minimises variations in cell density across the culture surface. Minimising such variations reduces the risk of variable outcomes which arise from variable cell density. By ensuring that cells are seeded in a similar manner irrespective of technique or user, it is less likely that results are simply an artefact of variable cell density. The UCS device is a simple to fabricate and use approach for standardisation of cell seeding density across users and experiments.

## Materials and Methods

We have used two methods to manufacture the devices used in the experiments. A simple and flexible route is the use of 3D printing which is becoming ubiquitous to most labs and research facilities. 3D printing exists in different formats with varying level of resolution, ranging from filament printing (FDM) to stereolithography (SLA). Filament printing is the most common 3D printing method but is also limited in resolution – in particular edge definition. 3D printing provides ease of device design to fit a wide range of applications from custom made samples to petri dishes and well plates (**Fig S1)**. The UCS designs were either made in SolidWorks or Rhino3D. STL files for some of the most common formats are provided as supplementary files. The devices are designed to provide a well-defined and constant separation distance between the surface and the UCS. This distance is provided by three legs placed at the periphery of the device. Near the edge, a 5mm inlet is provided to match a pipette tip through which the cell suspension is delivered. With the aim that the UCS could be widely adopted by the scientific community, we have also demonstrated volume scaling by injection moulding. Here, an insert was CNC milled in aluminium and fitted in the tooling of our injection moulding machine (Engel Victory 28 tons). UCS devices were then moulded in polycarbonate (Makrolon OD2015) with a cycle time of 10s. Such devices also benefit from full optical clarity which eases use during filling. Before use, UCS devices were either sterilised in 70% Ethanol for at least 1 hour or by UV irradiation. An alternative route for the injection moulded devices was autoclaving as the polycarbonate can sustain such a temperature treatment without failing (Tg 145°C).

### Measurement of cell settle time during UCS use

Settle time of cells delivered to a surface using the seeding device was assessed using a modified internal reflection microscopy setup. A 10mm thick glass plate was mounted on an inverted microscope, with LEDs used to illuminate through the glass plate in total internal reflection (TIR). As cells sediment to the surface, light is coupled out of TIR and imaged by a 4x objective placed perpendicular.

### hESCs pancreatic endoderm differentiation

Human embryonic stem cells (hESCs) (SA121, Cellartis AB) were cultured in the feeder-free and defined DEF-CSTM 500 system according to instruction from the supplier (Cellartis AB). The hESCs were cultured and differentiated in a humidified incubator with 5% CO2. For the differentiation towards pancreatic endoderm the following protocol was used. The cells were first differentiated to definitive endoderm in fibronectin coated cell culture flask using a highly effective patented protocol (WO 2012175633 A1). Subsequently, the cells were rinsed with PBS and dissociated to single cell suspensions with TrypLE Select at room temperature. Seeding of the cells was performed with 1000-3000 cells/mm2 in basal media with 100 ng/ml Activin A and 1 µM RockI.

The cells were seeded in 12 well plates using differentiation medium with 2% B27 at 3 different seeding densities both with and without the UCS device. With standard seeding, the wells were filled with 2ml of cell suspension. With the controlled seeding 0.3ml of cell suspension was injected into the UCS device and placed in the incubator. The UCS device was removed 1.5-2 hours after seeding and media was added to a total volume of 2ml per well. The pancreatic endoderm differentiation was initiated the day after seeding. The cells were differentiated towards pancreatic endoderm for 13 days with RPMI medium containing 64ng/ml FGF2 (Peprotech), 12% knockout serum replacement (Gibco) and 0.1% pen strep 21. Media was changed daily.

The cells were fixed by washing once with phosphate buffered saline (PBS) (Invitrogen and adding 4% formaldehyde (Lilly’s fixative, Mallinsckridt Baker) for 20 minutes. The cells were rinsed 2 times with PBS and permeabilized in 0.5% TritronX-100 (Sigma Aldrich) in PBS. The cells was washed in PBS and blocked with TNB blocking buffer (0.1M tris-HCL pH 7.5, 0.15M NaCl and 0.5% Blocking reagent from Perkin Almer TSA kit) for 30 minutes. Primary antibodies were diluted in in 0.1% TritronX- 100 in PBS and applied to the cells, followed by incubation overnight at 4C. The following primary antibodies were used: goat polyclonal anti-PDX1 (1:8000, Abcam, ab47383) and mouse anti-Nkx6.1 (1:500, in house facility, F55A10). Cells were washed 3 times with PBS for 5 minutes. The secondary antibodies Alexa Fluor 594 conjugated donkey-anti-mouse IgG (Invitrogen) and Alexa Fluor 488 donkey-anti-goat IgG (Invitrogen) were added in a 1:1000 dilution together with DAPI (1:2000) (SigmaAldrich) in 1% TritronX-100 in PBS for 45 minutes at room temperature. The cells were rinsed 3 times with PBS for 5 minutes.

### Image acquisition and quantification

Images were acquired using either an Olympus CX51 upright microscope equipped with a Prior motorized stage and 10X objective operated by ImageProPlus (Media Cybernetics, UK), or a GE InCell analyser with a 10X objective. Contiguous arrays were captured across the full extent of each plate well, including the curved wall which was later cropped from the image to leave only the complete culture area for analysis. Images were analysed using CellProfiler (Broad Institute, Harvard) to detect individual cells using DNA stains. After detection of individual cells, their centroids were computed. Processing of 1,000 2 megapixel images takes approximately 12h using CellProfiler 2.0 on an Intel Core i7 2600 CPU @ 2.4 GHz with 16Gb DDR2 RAM. The processing pipeline used in this work has been made available online at www.cellprofiler.org.

## Author Contributions and Notes

PMR, MOR, CHR, NG and MD designed research, PMR and CHR performed research, PMR analysed data; PMR wrote the paper, PMR, MOR, CHR, NG and MD edited the paper. This device was patented (application number PCT/GB2013/052776, publication number WO2O14O64449 A1; inventors: PMR, MOR, NG).

## Acknowledgments

Staff and technicians of the Centre for Cell Engineering and in the School of Biomedical Engineering at the University of Glasgow. The staff of the mechanical workshop in the School of Engineering at the University of Glasgow for their assistance in fabricating injection moulding tooling. We gratefully acknowledge test users of the device, and Marie Francene Cutiongco for necessary critical reading of the manuscript and valuable ideas to improve it. The work has been partially funded by the Engineering and Physical Sciences Research Council (EPSRC), Grant EP/F500424/1 DTC in Cell and Proteomic Technologies, and the Biotechnology and Biological Sciences Research Council (BBSRC) grant BB/K011235/1. C.H.R. and M.D. acknowledge financial support from Innovation Fund Denmark and Novo Nordisk A/S, Denmark.

## Data Availability and Material sharing

Designs for 3D printed UCS devices in various geometries are <u>available online</u> or by request. Injection moulded UCS devices may be available on request from the authors.

## Supplementary

**Figure S1:**
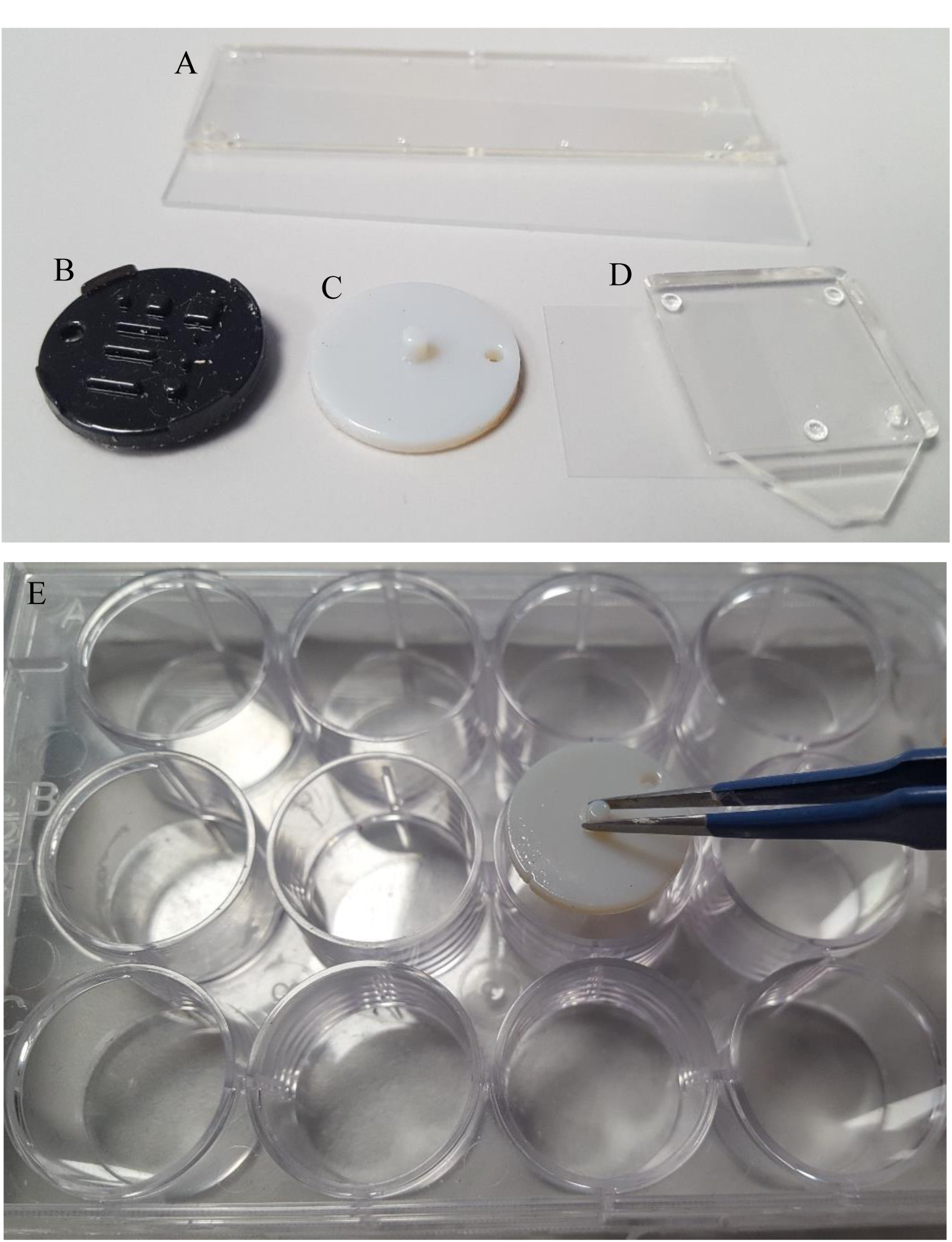
Various UCS geometries fabricated by injection moulding (A & D) and 3D printing (B & C). Parts A and D are used to seed cells uniformly on microscope slides and 22mm coverslips respectively. Part B allows for millimetre scale patterning with a specific design in a 12 well dish, whilst part C is used for uniform seeding in a 12 well dish, as shown in E.

**Figure S2:**
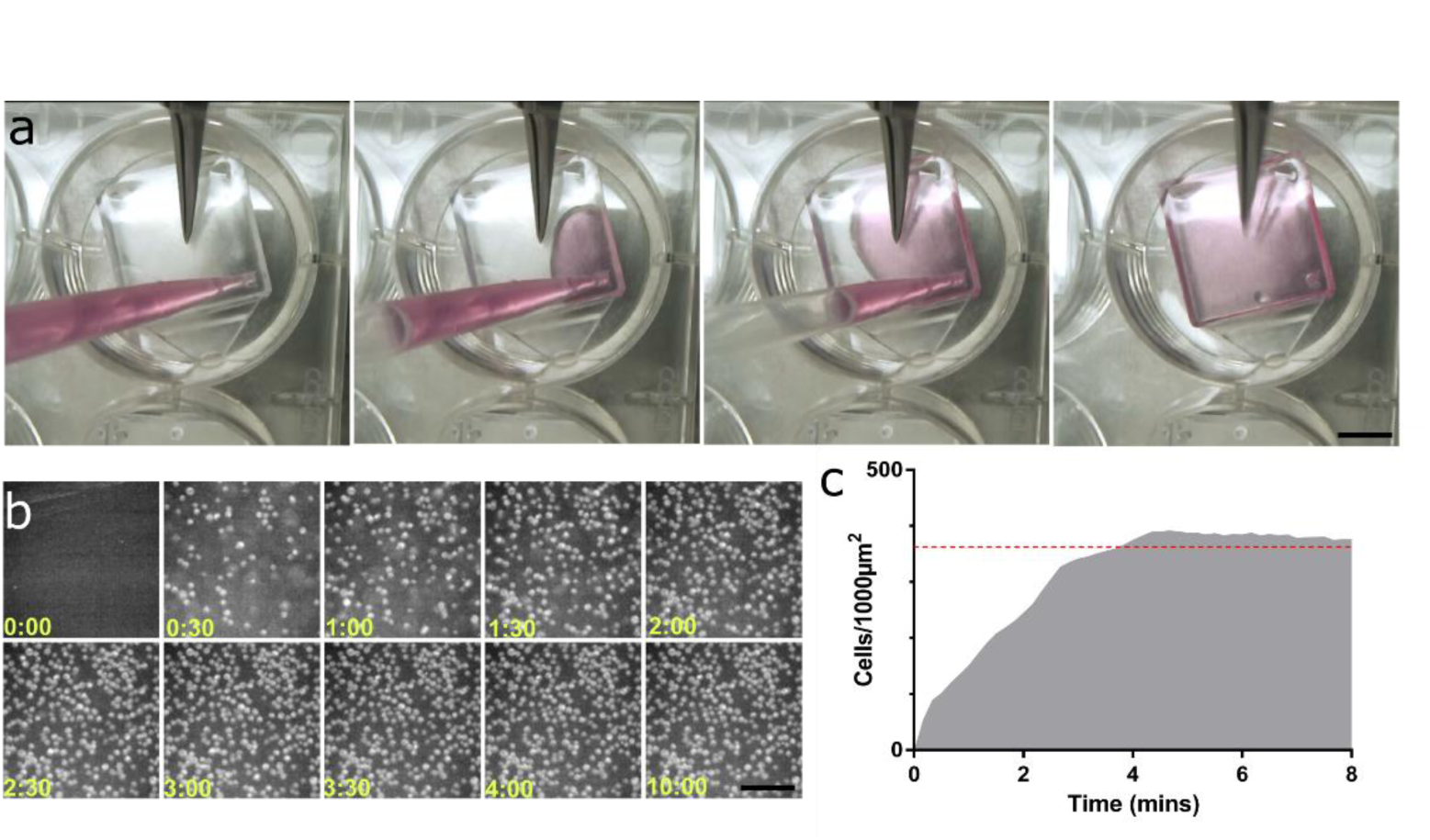
A) Cell suspension is added to the UCS device in under 3s. B) A modified internal reflection microscope was used to image cells as they settle to the culture surface over 10 mins. C) Target cell density is reached within 3 minutes at the target cell density (red dashed line).

**Figure S3:**
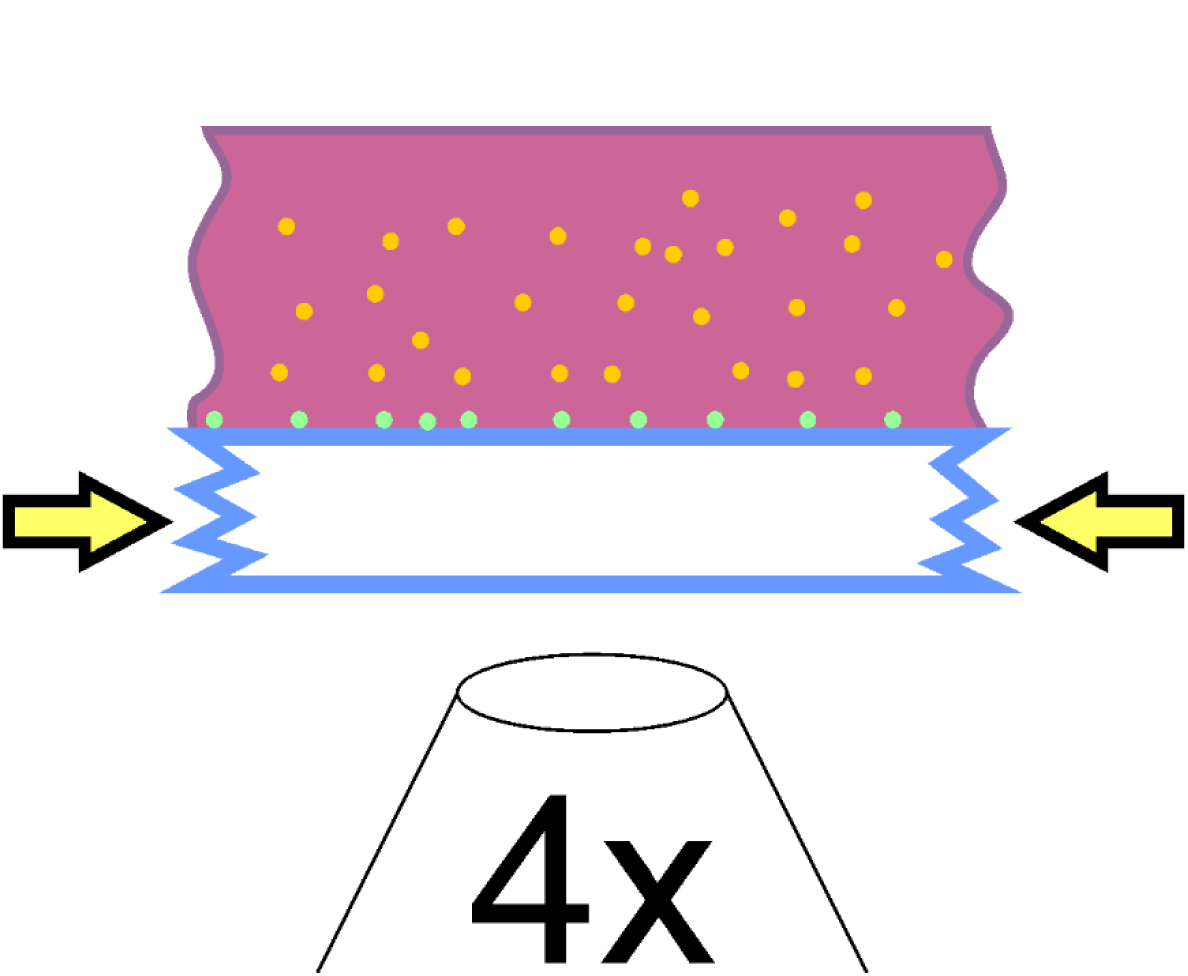
Schematic of modified internal reflection microscopy setup used to image cell sedimentation onto the culture surface in a wide field of view. LED illumination was coupled into a 4mm thick glass plate from the side, and cells seeded using the UCS device above a low magnification (4x) objective. The only illumination source was the side coupled illumination. As cells settle on the surface, light is coupled from the glass plate and collected by a camera. As shown in Fig S2 B.

**Figure S4:**
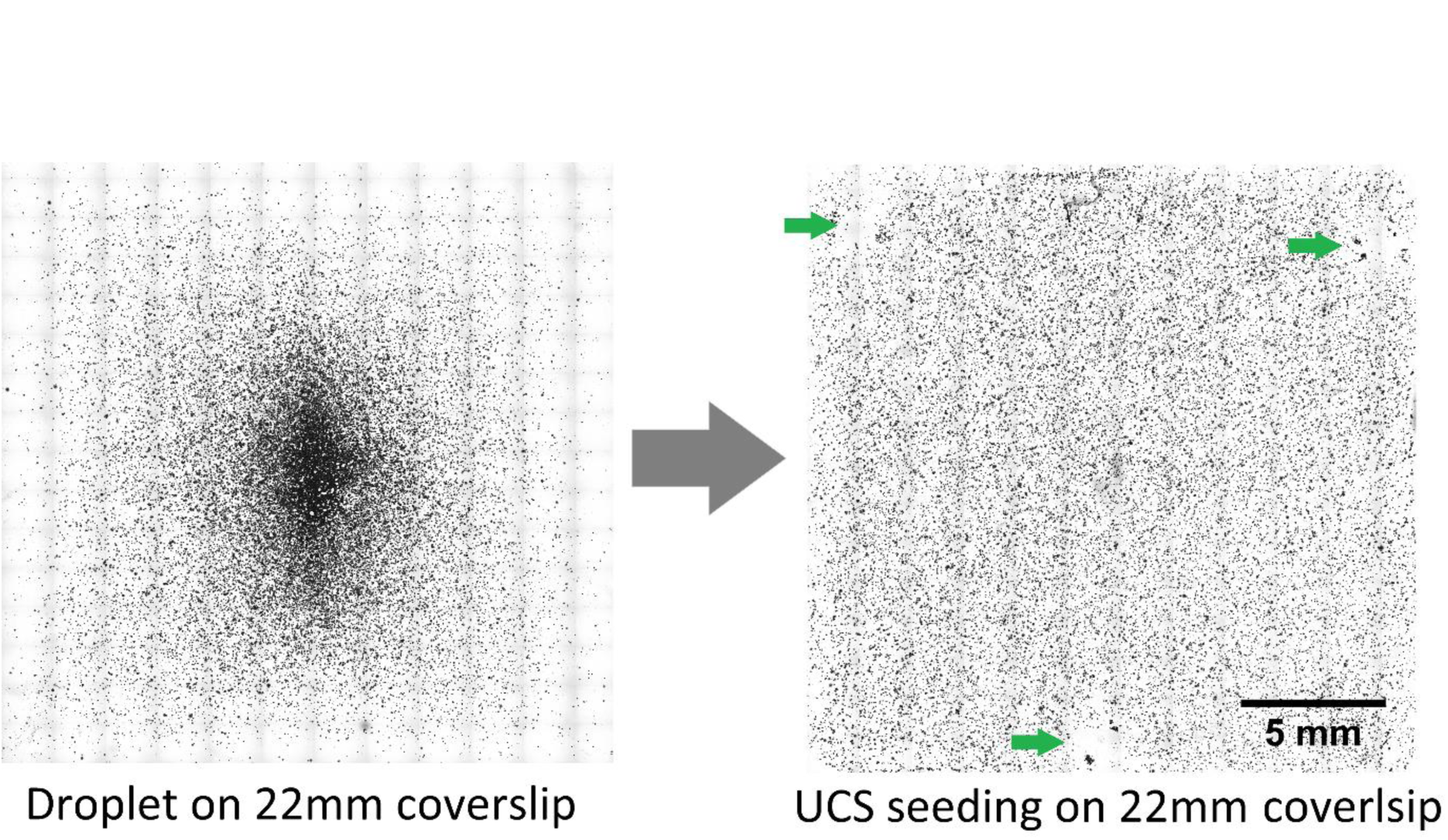
Correcting seeding variation on 22mm coverslip with the UCS device. Standard practice to add a droplet of cell suspension to the prepared coverslip creates a dome of fluid which distributes cells in an uneven manner. The UCS device (right) corrects this by confining the seeding suspension at a uniform height across the coverslip. Defects still occur at the location of spacers on the UCS device (green arrows) however they represent a much lower fraction of the culture area.

**Figure S5:**
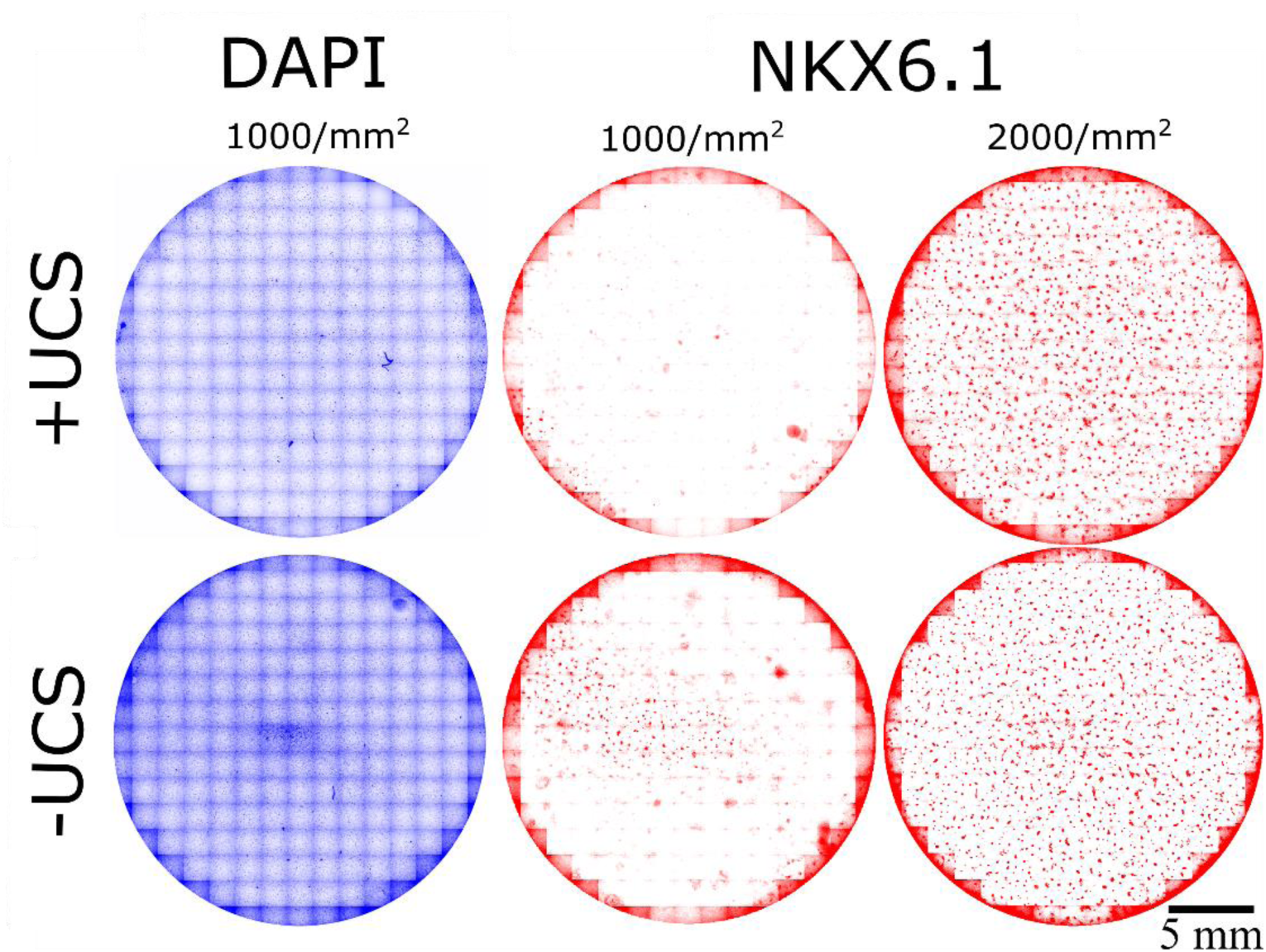
Uneven seeding at low densities may be beneficial. hESCs re-seeded at high plating densities before differentiation towards PE. At low density aggregation of cells in the well centre when seeded without the UCS provides a region of sufficient local density to stimulate differentiation, as evidenced by large clusters of Nkx6.1 positive cells. When variation in cell density is corrected using the UCS device there are reduced numbers of differentiated clusters. At higher densities of 2000/mm^2^ there is a more uniform cluster morphology across the dish when cells are seeded using the UCS. DAPI = blue, PDX6.1 = red.

## UCS device for contact-free wound models

Given that the uniformity of the cell density is given by the height of the confined gap, it is possible to take advantage of this to pattern cells during the seeding phase. If a device has varying heights it would be possible to adjust the cell density locally and create contact-free wound healing models, Fig. S6. Generally wound models are either performed as a physical injury by scraping or as an adhesive barrier during the early stages of cell culture. However, both are likely to damage the surface and may disrupt chemical or biological coatings important for the wound study. To demonstrate this effect we 3D printed a number of different designs using different 3D printers. Depending on the type of printer, different degrees of quality can be obtained. Filament printing (FDM) is the most common type of printer in a research environment. This type of printer has limited edge resolution and will thus give rise to rounded corners and consequently less defined wounds. On the other hand, stereolithography (SLA) printers can achieve much high edge resolutions and consequently the definition of the wound is much sharper. As a consequence, when making UCS’ for wound healing experiments, the wound definition is limited in resolution.

**Figure S6:**
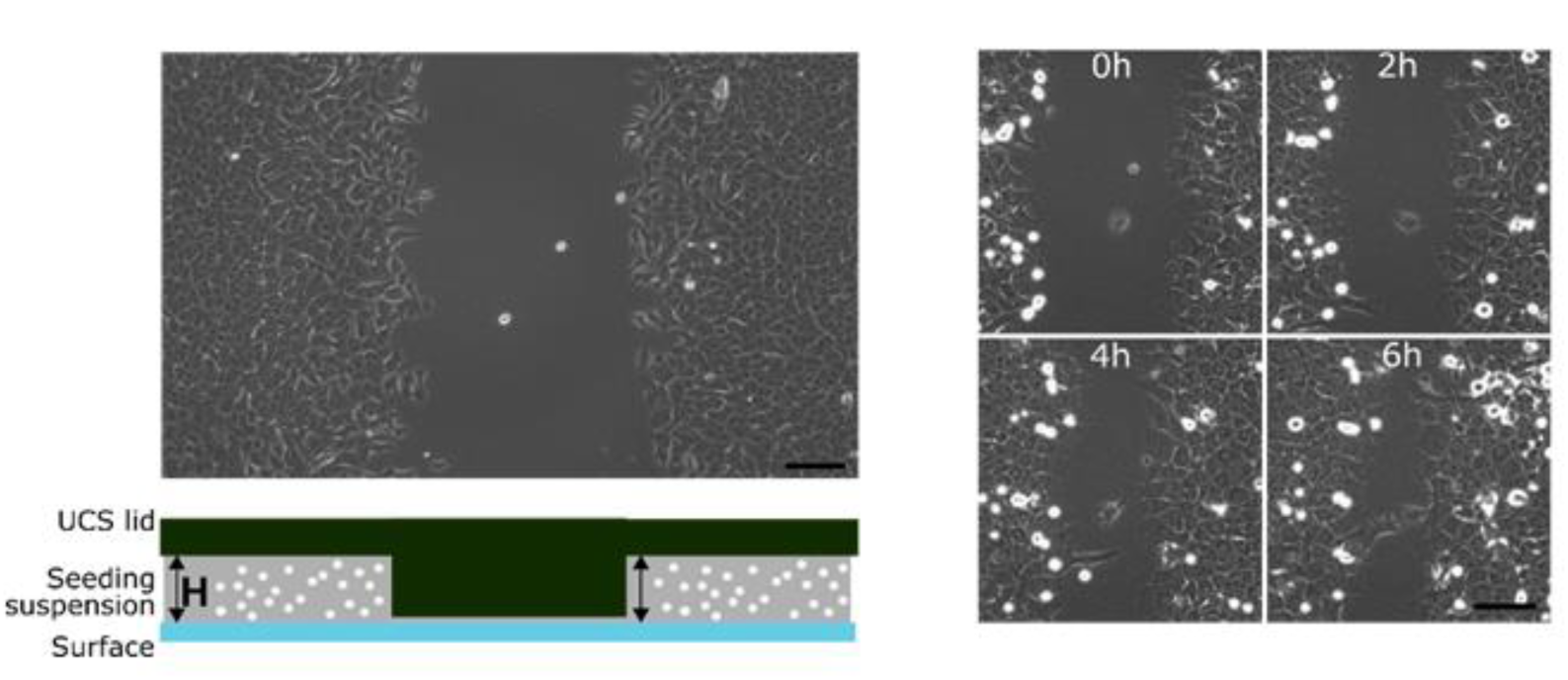
Contact-free gap closure assays created using the UCS. Variation of the distance between the UCS lid and the culture surface can effectively exclude cells from a region (left). After removal of the device, gap closure can be measured.

